# Transcranial Focused Ultrasound of the Human Subgenual Anterior Cingulate Reconfigures Resting State Connectivity: A Sham-Controlled fMRI Study

**DOI:** 10.64898/2025.12.10.693481

**Authors:** Amilcar Malave, Emily Valera, Andrew Birnbaum, Elisa Konofagou, Jacek P. Dmochowski

## Abstract

Transcranial focused ultrasound stimulation (tFUS) can modulate human brain activity, but its impact on large-scale functional connectivity (FC) *in vivo* remains incompletely characterized. Here we asked whether brief tFUS to the subgenual anterior cingulate cortex (sgACC) modulates its FC with other brain regions. In a sham-controlled, within-subject study of healthy adults (*N* = 16), we measured resting-state blood-oxygen-level dependent (BOLD) activity with functional magnetic resonance imaging (fMRI) before, during, and after five minutes of tFUS (i.e., five 20-second sonications followed by 40 second pauses). Subject-level sgACC–whole-brain FC was analyzed with a linear mixed-effects model (fixed: time, condition, time×condition; random: subject). tFUS increased sgACC-whole brain connectivity after sonication relative to sham. A smaller, non-significant trend was observed during sonication. Baseline-controlled analyses were employed to rule out the possibility that the observed increase in FC was due to a “regression to the mean” effect. Interestingly, network-level analysis revealed a clear disassociation in the evolution of resting-state FC across the 15 minute recordings: during sham sessions, the sgACC’s connectivity with the default mode network (DMN) increased, while FC with other networks was largely unchanged. On the other hand, active tFUS led to a notable increase in FC with non-DMN networks (especially the cognitive control network), while connectivity with the DMN was stable. These results provide whole-brain evidence that tFUS reconfigures network dynamics in the human brain, motivating clinical applications to disorders characterized by dysregulated brain activity.

## Introduction

Transcranial focused ultrasound stimulation (tFUS) has emerged as a noninvasive neuromodulation approach with millimeter-scale focality and the ability to access subcortical and medial structures that are challenging to reach with electromagnetic methods such as transcranial magnetic stimulation (TMS) or transcranial direct current stimulation (tDCS) (1, 2). Foundational studies spanning *in vitro*, small-animal, and human preparations established feasibility and provided early demonstrations of state-dependent effects (3–5). These advances have catalyzed a growing body of research applying tFUS to probe and perturb human brain networks (6–13).

Despite this progress, the biophysical mechanisms by which low-intensity ultrasound influences neural activity remain an active area of investigation (14–16). Proposed accounts include direct effects on mechanosensitive ion channels (17– 19), modulation of membrane capacitance and bilayer elasticity consistent with intramembrane cavitation (20, 21), flexoelectric coupling of lipid membranes (22), and indirect astrocytic or neurovascular contributions (23). Clarifying these mechanisms is essential for developing principled strategies to bias excitation–inhibition balance and to achieve reproducible, directionally specific changes in circuit activity.

Concurrently, there is growing interest in how tFUS alters large-scale functional organization of the human brain. Functional connectivity (FC) – typically operationalized as correlations in the blood-oxygen-level–dependent (BOLD) signal between regions – provides a macroscopic readout that is sensitive to behavioral state and clinical phenotype (24, 25). Human functional magnetic resonance imaging (fMRI) studies have reported both increases and decreases in FC following tFUS, likely reflecting differences in targeted circuits, sonication parameters, behavioral context, and analysis choices (10, 12, 13, 26–30). A consistent picture has yet to emerge regarding when and where tFUS strengthens versus weakens coupling.

Here we focus on the subgenual anterior cingulate cortex (sgACC), a deep medial prefrontal node implicated in affect regulation and prominently linked to mood disorders (31, 32). We conducted a within-subject, randomized, sham-controlled study in healthy adults (two sessions per participant) combining tFUS with concurrent fMRI. Our primary endpoint was BOLD FC between the sgACC and the rest of the brain, estimated at the subject level and analyzed with linear mixedeffects models.

We found that: (i) sgACC–whole-brain FC increased from pre- to post-sonication in Active relative to Sham (time × condition Pre → Post contrast), with a similar but non-significant trend during tFUS, (ii) this effect was not explained by regression-to-the-mean or baseline differences, as it persisted in baseline-controlled models, and (iii) most importantly, the added coupling was expressed selectively across canonical brain networks: Sham sessions showed an increase in sgACC coupling with the default mode network (DMN) over time, whereas Active tFUS preferentially increased sgACC coupling with non-DMN systems, most notably the Cognitive Control network, during and following sonication.’’

## Materials and Methods

### Participants

Sixteen healthy adults (7 females; age 26*±* 7 years, mean *±* standard deviation) participated in a within-subjects study consisting of two sessions employing Active and Sham tFUS. Sessions were spaced two weeks apart to ensure sufficient washout of effects. The order of the sessions was randomized and counterbalanced across the cohort. Experimental procedures were approved by the Institutional Review Board of the City University of New York, and subjects provided written informed consent prior to participation.

### tFUS Parameters

tFUS was delivered at 500 kHz with a circular transducer (Sonic Concepts CTX-500) driven by an integrated function generator and amplifier (Sonic Concepts NeuroFUS; Fig 1a). The waveform was pulsed with a repetition frequency (PRF) of 40 Hz and a duty cycle of 30%. Sonication duration was 20 s, with a 40 s interval between successive sonications (Fig 1e). All subjects received a total of 5 sonications. Acoustic intensity was measured in a water tank under free-field conditions with a calibrated hydrophone (ONDA corporation) as *I*_spta_ = 667 mW/cm^2^. The empirical beampattern is shown in Fig 1c.

**Fig. 1.**
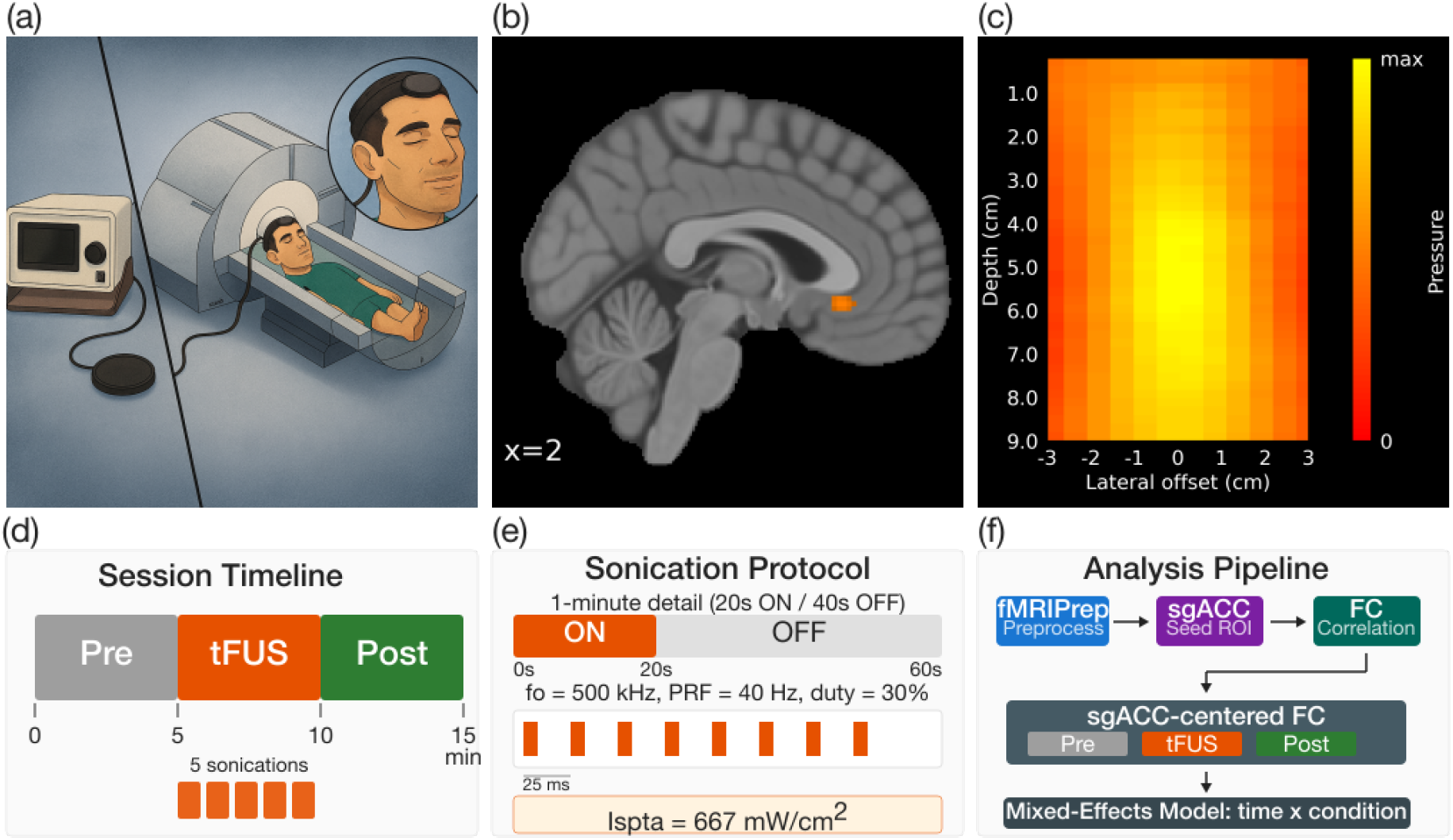
Experimental overview. **(a)** MRI-compatible setup for transcranial focused ultrasound stimulation (tFUS) during 3 T resting-state fMRI. A circular transducer is secured to the forehead with an elastic headband to target the (right) subgenual anterior cingulate cortex (sgACC). Participants (*N* = 16) completed two counterbalanced sessions (Active, Sham) spaced two weeks apart. **(b)** Sagittal MRI slice highlighting the location of the sgACC. **(c)** Empirical beampattern measured in a water tank (500 kHz carrier) showing lateral spread and focal depth (i.e., approximately 6 cm); free-field spatial-peak temporal-average intensity *I*_spta_ = 667 mW*/*cm^2^. **(d)** Session timeline: a 15-min BOLD acquisition was divided into three 5-min windows (Pre, tFUS, Post). In Active sessions, sonication occurred only in the middle window. **(e)** One-minute zoom of the stimulation train used during the tFUS window: five cycles of 20 s ON / 40 s OFF; within ON-epochs, pulses were delivered at 40 Hz and 30% duty. **(f)** Analysis pipeline: data were preprocessed with fMRIPrep; sgACC-centered functional connectivity (FC; Fisher-z correlations to all parcels) was computed per subject, condition, and window. The prespecified primary test was a subject-level linear mixed-effects model with fixed effects for time, condition, and their interaction, evaluating the Pre to Post difference-in-differences (DiD, Active-Sham, Post-Pre).

The transducer positioning relied on a computational procedure to identify the subject-specific optimal scalp location for targeting the sgACC (nominal target location shown in Fig 1b). Namely, the subject’s anatomical MRI was employed to exhaustively search across a broad range of discretized scalp positions to identify the coordinates on the forehead (frontal pole region) whose normal vector most closely intersected the right sgACC. This resulted in a variable distance to the target across subjects ranging from 5 to 6.5 cm. The focal depth of the transducer was set to 6 cm and verified empirically with measurements in a water tank (Fig 1c). The transducer was fixed to the subject’s scalp using an elastic headband (Fig 1a).

### MRI and fMRI

Imaging was performed at 3 T with a Siemens Magnetom Prisma. Structural images were acquired with a T1-weighted MPRAGE sequence employing isotropic 0.8 mm voxels. Resting-state functional BOLD scans were acquired with an echo-planar imaging (EPI) sequence employing 2 mm isotropic voxels, a repetition time (TR) of 1000 ms, echo time (TE) of 37 ms, a flip angle of 52 degrees, and posterior–to–anterior phase encoding. Subjects were instructed to rest but stay awake and to not think about anything in particular.

The duration of the BOLD acquisition was 15 minutes (900 TRs). During the Active tFUS session, sonication commenced at 300 s (Fig 1d); no sonication was applied during the Sham session. Throughout the manuscript, we refer to the 5 minute pre-stimulation time window as “baseline” or “Pre”. Similarily, we denote the last 5 minutes of each BOLD acquisition as “Post”.

Subjects did not report perceptions of tFUS after the conclusion of their two sessions.

### Anatomical preprocessing

Processing of anatomical images was performed in *fMRIPrep* (33–36) (Fig 1f). T1-weighted (T1w) images were corrected for intensity non-uniformity (INU) with `N4BiasFieldCorrection` (37), distributed with ANTs (38), and used as the T1w-reference throughout the workflow. The T1w-reference was then skull-stripped with a *Nipype* implementation of the `antsBrainExtraction.sh`workflow (from ANTs), using OASIS30ANTs as the target template. Brain tissue segmentation of cerebrospinal fluid (CSF), white-matter (WM) and gray-matter (GM) was performed on the brain-extracted T1w using `fast`(39). Brain surfaces were reconstructed using `recon-all`(40), and the brain mask estimated previously was refined with a custom variation of the method to reconcile ANTs-derived and FreeSurfer-derived segmentations of the cortical gray-matter of Mindboggle (41). Volume-based spatial normalization to a standard space – the ICBM 152 Nonlinear Asymmetrical template, version 2009c (42) – was performed through nonlinear registration with `antsRegistration`(ANTs 2.5.1), using brain-extracted versions of both the T1w reference and the T1w template.

### BOLD preprocessing

All functional MRI data were preprocessed with `fMRIPrep`(v24.0.0) (33, 34). For each run, fMRIPrep applied slice–timing correction, estimated head motion relative to a reference volume with MCFLIRT (43), and coregistered the BOLD reference to each participant’s T1-weighted anatomy using FreeSurfer’s boundary–based registration (BBR; 6 degrees of freedom) (44). Outputs were written to the MNI152NLin2009cAsym space along with confound time series (e.g., framewise displacement, DVARS, CompCor, global signals) and HTML reports for quality control.

Subsequent denoising and time–series extraction were performed in Python with *Nilearn* (45) using a probabilistic functional parcellation. Specifically, we used the `DiFuMo`(Dictionary of Functional Modes) probabilistic atlas (46). Because this atlas provides probabilistic component maps rather than hard labels, we employed the Nilearn function `NiftiMapsMasker`to project the atlas maps onto each preprocessed BOLD run. Regional time series were then standardized (z–scored) and band–pass filtered (0.01–0.10 Hz). We regressed out the six rigid–body motion parameters (translations and rotations) and their first temporal derivatives.

In summary, preprocessing followed a standard fMRIPrep workflow (slice–timing correction, motion estimation, BBR to T1w, MNI normalization) and custom Python code then extracted probabilistic–atlas regional signals with minimal motion regression and 0.01–0.10 Hz filtering to produce condition–specific time series used in all downstream connectivity analyses.

### Whole-brain and network-level effects

Our inferential plan followed a hierarchical design. As the primary analysis, we asked whether the temporal trajectory of sgACC FC differed between Active and Sham conditions. Let FC_*ict*_ denote the subject-level mean sgACC–whole-brain FC for subject *i*, condition *c* ∈ {Sham, Active}, and time window *t* ∈ {Pre, tFUS, Post}. We fit a linear mixed-effects model with treatment coding (reference values of *Sham* and *Pre* for condition and time window, respectively), fixed effects for time, condition, and their interaction, and a random intercept for subject:

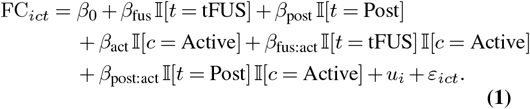

where I[·] is an indicator function, *β*_0_ models the mean FC of the Sham baseline, *β*_fus_ and *β*_post_ are within-sham changes versus Pre, *β*_act_ is the Active–Sham difference at Pre, and the interaction terms are difference-in-differences (DiD) contrasts: *β*_fus:act_ tests whether the Pre → tFUS change differs between Active and Sham, and *β*_post:act_ tests whether the Pre → Post change differs between Active and Sham. The random intercept 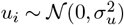 captures subject-specific baselines; *ε*_*ict*_ ∼ 𝒩 (0, *σ*^2^) are the model residuals. The prespecified contrast was the two-sided test of the Pre → Post DiD (*H*_0_: *β*_post:act_ = 0, *α* = 0.05). A *planned* secondary contrast evaluated the peri-stimulation DiD (*H*_0_: *β*_fus:act_ = 0). To avoid treating edges as independent, FC was aggregated to the subject level within each (condition, time) cell prior to modeling; inference on fixed effects used Wald *z*-tests.

Conditioned on observing a significant *positive* whole-brain effect in the primary test, we conducted secondary networklevel analyses to determine which large-scale networks absorbed the additional sgACC coupling. For each canonical brain network in the Yeo-7 network parcellation (47) (Visual, Cognitive Control, Default Mode, Dorsal Attention, Limbic, Salience, Somatomotor), we formed subject-level baseline-referenced changes (ΔFC) and tested the directional hypothesis that Active stimulation increases sgACC connectivity more than Sham:

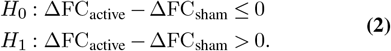

These were implemented as subject-level paired t-tests comparing baseline-referenced ΔFC between Active and Sham conditions. Multiple comparisons across networks and time windows were controlled using the Benjamini–Hochberg false discovery rate at *q* = 0.05. For interpretability, we report effect sizes (mean differences in ΔFC) with two-sided 95% confidence intervals alongside the one-sided *p* -values. All tests, sidedness, and error-rate control procedures were specified *a priori* at the analysis-family level as described above.

### Analysis of baseline FC effects

To assess whether the observed connectivity changes could be attributed to a “regression-to-the-mean” (RTM), we examined how baseline sgACC connectivity relates to the subsequent change, ΔFC, and whether the relationship between baseline FC and ΔFC differs between conditions. For each subject *i*, condition *c* ∈ {Sham, Active}, and follow-up window *t* ∈ {tFUS, Post}, we defined:

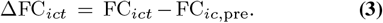

We then fit an ordinary least squares (OLS) model that pooled tFUS and Post windows for each subject × condition (two rows per subject × condition; *N* = 16 × 2 × 2 = 64). The model uses subject fixed effects (*α*_*i*_) and a time-window fixed effect (with tFUS as the reference) so that mean differences between tFUS and Post are absorbed without estimating separate slopes per window:

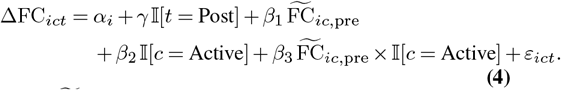

Here, 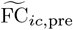 is the centered subject × condition baseline. The coefficient *γ* captures the post–tFUS ΔFC that is *shared* across conditions. The primary hypothesis test is *H*_0_: *β*_3_ = 0 (i.e., equivalent regression slopes in Sham and Active). Inference used cluster-robust standard errors with subjects as clusters. For visualization (Fig. 3), we plotted baseline FC vs. ΔFC separately for Sham and Active conditions, displaying window-specific markers and the corresponding regression line from the OLS model.

### Distance as a moderator of the time × condition effect

We tested whether scalp-to-sgACC distance moderates the effect of tFUS on sgACC-centered FC. For each subject *i*, condition *c* ∈ {Sham, Active}, and time window *t* ∈ {Pre, tFUS, Post}, we fit a linear mixed-effects model with a subject-specific random intercept:

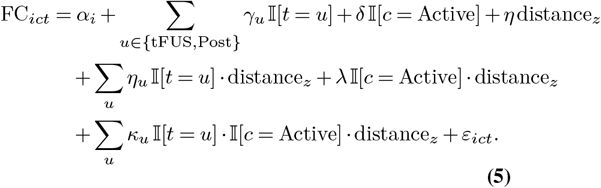

where *α*_*i*_ is a subject-specific random intercept, *γ*_*u*_, *u* ∈ {tFUS, Post} are fixed effects for time-window, *η* is a fixed effect for z-scored scalp-to-sgACC distance (distance_*z*_, mean = 58 mm, SD = 5 mm), *η*_*u*_ and *λ* capture the interactions between time-distance and condition-distance, respectively, and finally, *κ*_*u*_ is the three-way interaction between time window, condition, and distance.

Treatment coding used Pre (time) and Sham (condition) as references. The moderation analysis tests for a significant effect of distance on the time-condition interaction: *H*_0_: *κ*_post_ = 0.

In order to compute the scalp-to-sgACC distance, we manually measured the distance of the line segment connecting the center of the transducer to the visually identified sgACC location on the T1w MRI.

### Distance analysis: difference-in-differences correlation

To visualize the relationship between connectivity change and scalp-to-sgACC distance, we correlated each participant’s scalp-to-sgACC distance to the DiD of the sgACC coupling change. For subject *i* and time window *t* ∈ {tFUS, Post} we computed baseline-referenced changes:

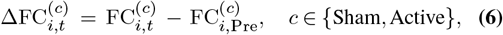

and the paired DiD contrast (Eq. Eq. (7)):

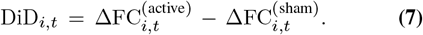

In Figure 5, we plot the scatter and report Spearman correlation coefficients *ρ* between D iD_*i,t*_ and scalp-to-sgACC distance (mm). We report two-sided 95% CIs via 10,000 bootstraped resamples, together with slope estimates computed with both OLS and Theil–Sen robust regression (units of ΔFC per 10 mm).

## Results

### Increased sgACC-whole-brain connectivity after Active tFUS

To formally test whether tFUS modulated sgACC-centered connectivity beyond Sham, we fit a subject-level linear mixed-effects model. For each subject, condition (Sham, Active), and time window (Pre, tFUS, Post), we computed the mean FC between the sgACC and all other atlas parcels (1023), yielding one sgACC–whole-brain FC value per subject-time-condition (16 subjects ×2 conditions ×3 time windows = 96 observations). Time window and condition were modeled as categorical fixed effects with *Pre* and *Sham* as reference levels, and a random intercept for subject accounted for repeated measures. This parameterization allowed us to directly test whether the change in sgACC FC from Pre to Post (and from Pre to tFUS) differed between Active and Sham sessions (time ×condition interaction).

The results of the model are presented in Table 1. Within the Sham condition, sgACC FC did not change reliably over time (tFUS vs. Pre: *β* = −0.020, *p* = 0.444; Post vs. Pre: *β* = −0.011, *p* = 0.671), indicating a lack of systematic drift in sgACC-centered FC. Importantly, the post-sonication time × condition interaction was significant: t he additional Pre-to-Post change in the Active session relative to Sham was positive (*β* = 0.079, 95% CI [0.006, 0.151], *p* = 0.033), indicating a significant enhancement o f s gACC F C. T he corresponding interaction at the tFUS window showed a similar but nonsignificant tendency (*β* = 0.060, *p* = 0.104). Together, these results support a conservative but robust conclusion that Active tFUS to the sgACC is associated with increased post-sonication connectivity.

**Table 1.** Linear mixed-effects model of subject-level mean sgACC FC. The dependent variable is the mean sgACC–whole-brain FC for each subject, condition, and time window. Time window (Pre, tFUS, Post) and condition (Sham, Active) are coded categorically with *Pre* and *Sham* as reference levels. The model includes a random interceptfor subject. The key effect of interest is the post-sonication time × condition interaction, indicating a larger Pre-to-Post increase in sgACC FC for Active tFUS relative to Sham.

| Effect | Estimate | SE | $z$ | $p$-value | 95% CI |
| --- | --- | --- | --- | --- | --- |
| Intercept (Sham, Pre) | 0.121 | 0.021 | 5.89 | < 0.001 | [0.081, 0.161] |
| Time: tFUS vs. Pre (Sham) | -0.020 | 0.026 | -0.77 | 0.444 | [-0.071, 0.031] |
| Time: Post vs. Pre (Sham) | -0.011 | 0.026 | -0.42 | 0.671 | [-0.062, 0.040] |
| Condition: Active vs. Sham (at Pre) | -0.061 | 0.026 | -2.34 | 0.019 | [-0.112, -0.010] |
| Interaction: (tFUS vs. Pre) × (Active vs. Sham) | 0.060 | 0.037 | 1.63 | 0.104 | [-0.012, 0.133] |
| **Interaction: (Post vs. Pre) × (Active vs. Sham)** | **0.079** | **0.037** | **2.13** | **0.033** | **[0.006, 0.151]** |
| Subject random intercept variance | 0.001 |  |  |  |  |
| Residual variance | 0.0055 |  |  |  |  |

These subject-level effects are visualized for Sham and Active sessions in Fig. 2. In the Sham condition (Fig. 2a), sgACC FC exhibits no systematic monotonic change from Pre to tFUS to Post, and individual trajectories fluctuate around a stable mean. In contrast, the Active tFUS condition (Fig. 2b) shows a clear “ramping” pattern: sgACC FC increases from a lower baseline to higher values during tFUS and reaches its highest levels post-sonication. The alignment of individual trajectories with the model-based interaction effect visually reinforces the conclusion that tFUS strengthens connectivity with the sgACC in a manner that outlasts the stimulation.

**Fig. 2.**
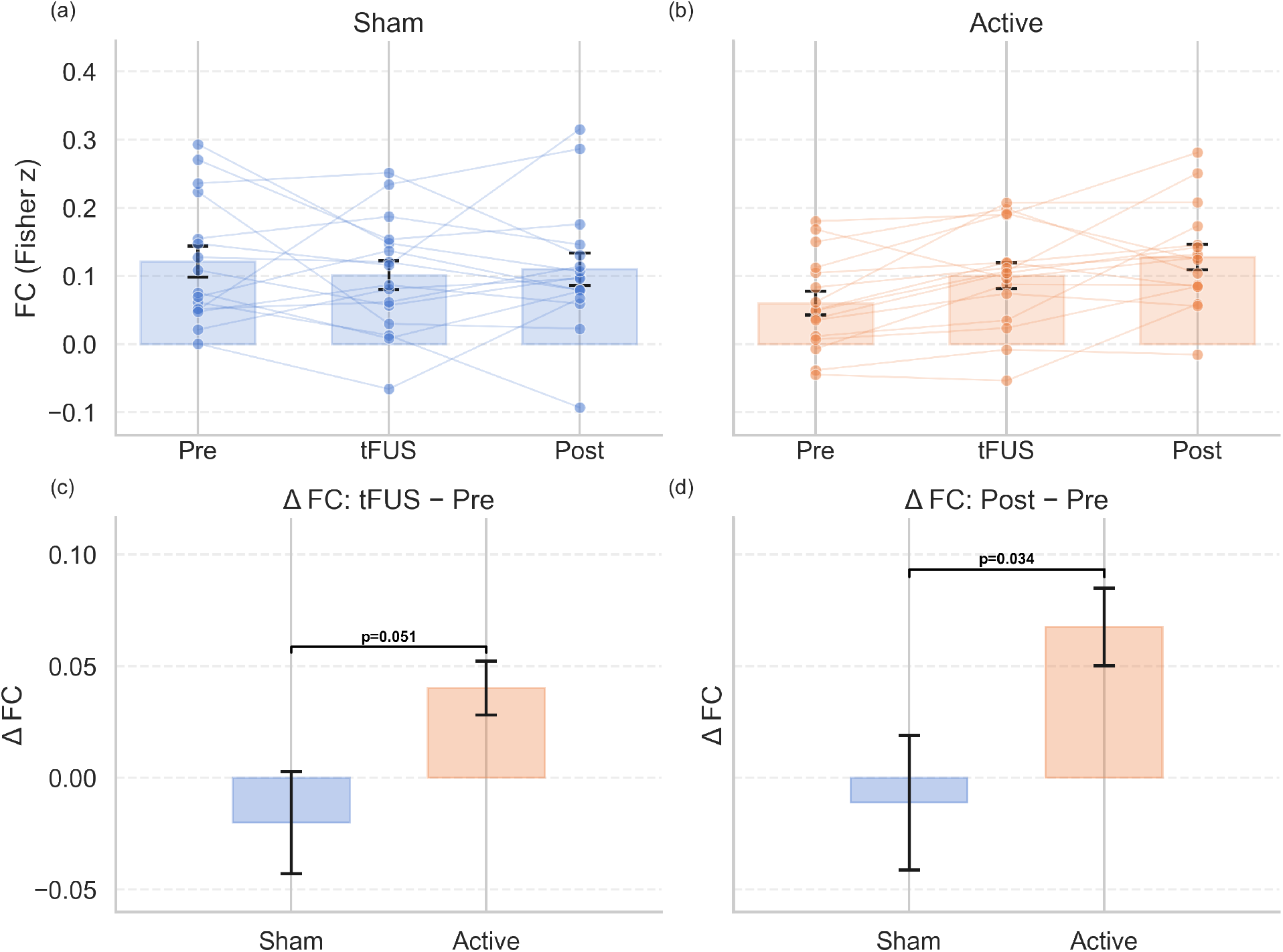
tFUS increases connectivity of the sgACC after Active but not Sham stimulation. **(a)** *Sham tFUS*. Scatter-overlaid bar plots show the distribution of sgACC–wholebrain FC at each window (Pre, tFUS, Post); markers denote subject means and thin lines connect subjects across time. FC remains stable across time, consistent with the results of the mixed-effects model (tFUS vs. pre: *β* = −0.020, *p* = 0.444; post vs. pre: *β* = 0.011, *p* = 0.671). **(b)** *Active tFUS*. The same visualization reveals a “ramping” trajectory from baseline through post-sonication. The difference-in-differences (Δ*FC*_Active_ − Δ*FC*_Sham_) is significantly positive after sonication (Active–Sham, Pre → Post: *β* = 0.079, 95% CI [0.006, 0.151], *p* = 0.033), indicating a stimulation-specific increase in sgACC FC. A similar, smaller trend is present during tFUS (Pre → tFUS interaction: *β* = 0.060, *p* = 0.104) **(c)** *Baseline-referenced connectivity change during tFUS*. Bars show subject-level means of ΔFC = FC_tFUS_ − FC_Pre_ (error bars depict 95% CI). A planned post-hoc contrast of Active versus Sham shows a trend towards a greater increase in FC during Active stimulation (Active–Sham ΔFC = +0.060, *p* = 0.051). **(d)** *Baseline-referenced connectivity change post-sonication*. Same as (c) but now shown for the post-sonication period: ΔFC = FC_Post_ − FC_Pre_. The corresponding post-hoc contrast is significant (Active–Sham ΔFC = +0.079, 95% CI [0.006, 0.151], *p* = 0.034).

**Fig. 3.**
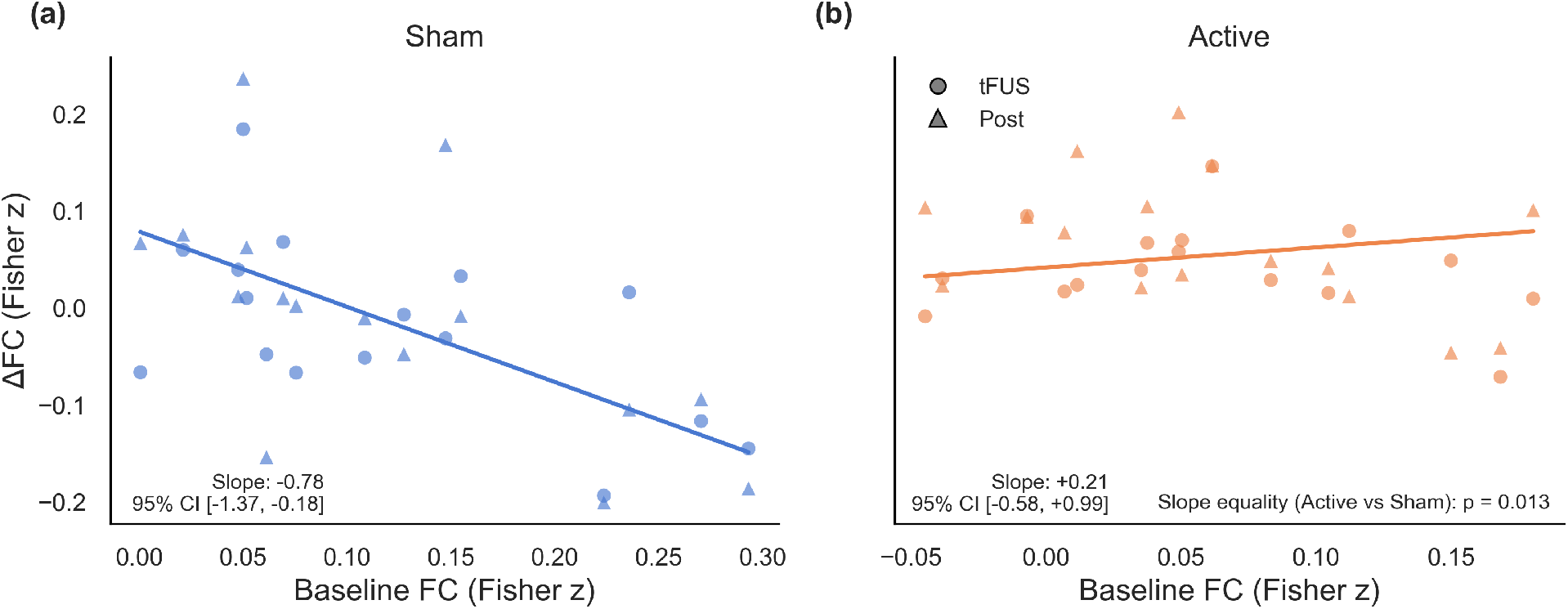
Baseline connectivity level predicts FC change in Sham but not Active tFUS. (a) Sham tFUS: subject-level scatter of baseline sgACC FC (x-axis) versus change from baseline ΔFC = FC_*t*_ − FCpre (y-axis), with data pooled across tFUS (º) and post-sonication (△) windows. The regression line is derived from an OLS model with fixed effects for subject, time-window, and condition. The inverse relationship between ΔFC and baseline FC observed in the Sham condition is consistent with a regression-to-the-mean account. (b) Same as (a) but now shown for Active tFUS. The slope of the regression line is mildly positive and does not differ significantly from zero. Importantly, the difference in slopes between Active and Sham is significant (*p* = 0.013), indicating that baseline effects are not driving the observed post-sonication connectivity increase during Active stimulation.

Panels c–d plot baseline-referenced changes (ΔFC = FC_*t*_ − FC_pre_) for the tFUS and post-sonication windows, respectively. Sham values remain centered near zero across windows, whereas the FC change with Active stimulation is positive both during and after stimulation. A planned post-hoc contrast of ΔFC between Active and Sham indicated a borderline significant trend during the tFUS window (Active–Sham ΔFC = +0.060; *p* = 0.051) and a significant effect post-sonication (Active–Sham ΔFC = +0.079; *p* = 0.034).

### Baseline connectivity level predicts FC change in Sham but not Active tFUS

Due to the different baseline FC values between Sham and Active sessions, it was important to rule out that discrepancies in baseline level drive the observed post-sonication connectivity increase (a “regression-to-the-mean” effect). To address this, we modeled the change from baseline (Δ*FC*) as the dependent variable using OLS with fixed effects for subject, time window, condition, and baseline FC, as well as an interaction term between baseline and condition.

The model explained substantial variance in ΔFC (*R*^2^ = 0.676; *p* = 0.017). In Sham, higher baseline sgACC FC predicted larger subsequent decreases in connectivity (slope = −0.776, 95% CI [−1.37, −0.18], *p* = 0.010; Table 2), consistent with a regression-to-the-mean account. In the Active condition, however, the baseline–change association did not significantly differ from zero (slope +0.207, 95% CI [−0.575, 0.989], n.s.). Critically, the slope difference between conditions was significant (*β*_3_ = +0.983, 95% CI [0.203, 1.763], *p* = 0.013), indicating that the mapping from baseline to subsequent change differs under stimulation. Figure 3 shows the relationship between baseline connectivity and subsequent FC change (circle markers denote the tFUS window, triangles denote the post-sonication window). The inverse relationship between baseline level and FC difference is evident for Sham (Panel a), while the slope of the regression line is mildly positive in the Active condition (Panel b).

**Table 2.**
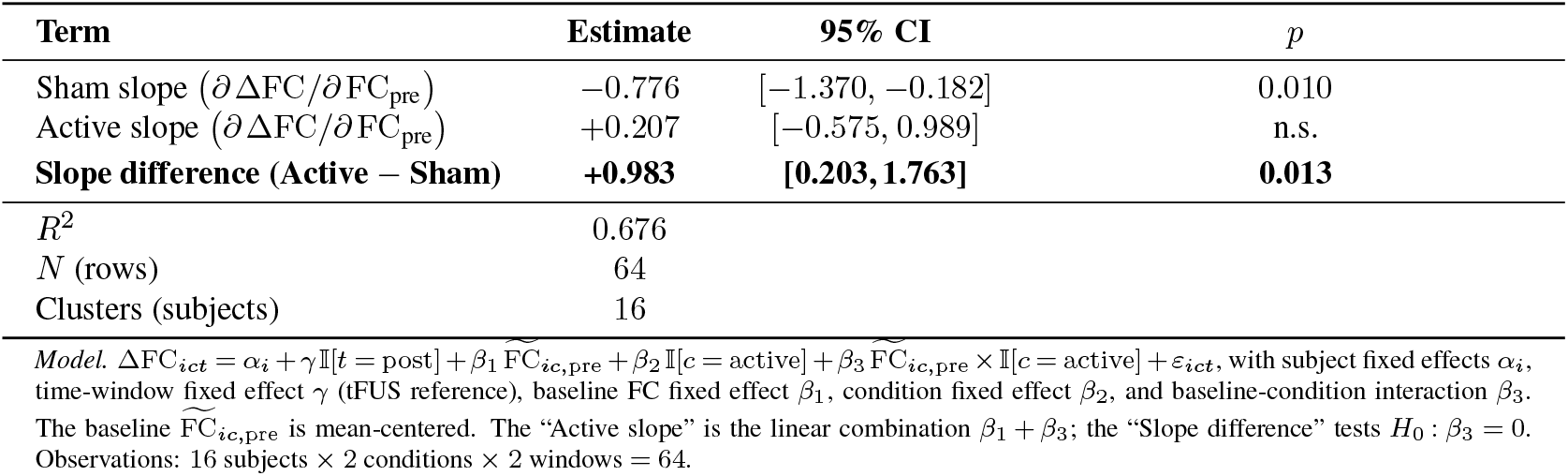
Testing the relationship between baseline FC and the subsequent change in connectivity (“Baseline–change slope test”). We fit an ordinary least squares model with fixed effects for subject, time-window (categorical with tFUS reference), condition (categorical with Sham reference), and baseline FC (z-scored). The dependent variable is ΔFC_*ict*_ = FC_*ict*_ − FC_*ic*,pre_ for subject *i*, condition *c* and time window *t*. We assay whether the baseline–change slopes are significantly different between Sham and Active.

### Network-level redistribution of sgACC connectivity

Having confirmed a post-sonication increase in sgACC connectivity under Active stimulation, we next examined which large-scale networks expressed this additional coupling. Using the Yeo-7 parcellation (47) (Visual, Cognitive Control, Default Mode, Dorsal Attention, Limbic, Salience, and Somatomotor networks), we compared subject-level baseline-referenced ΔFC at the tFUS and Post time windows (summarized in Table 3). A clear dissociation emerged: in *Sham*, increases from baseline were confined to the Default Mode

**Table 3.** Network-specific tests of Active–Sham differences in baseline-referenced ΔFC. One-sided hypotheses *H*_1_: ΔFC_active_ *>* ΔFC_sham_ were evaluated per network and window; *p* values are one-sided and FDR corrected within the family of network × window tests (*q*).

| Network | Window | $n_{\text{sham}}$ | $n_{\text{active}}$ | $t$ | $p$ | $q_{\text{FDR}}$ |
| --- | --- | --- | --- | --- | --- | --- |
| Visual | tFUS | 16 | 16 | 0.884 | 0.187 | 0.219 |
| Visual | Post | 16 | 16 | 1.129 | 0.132 | 0.193 |
| **Control** | **tFUS** | **16** | **16** | **3.139** | **0.0024** | **0.028** |
| **Control** | **Post** | **16** | **16** | **2.860** | **0.0040** | **0.028** |
| Default Mode | tFUS | 16 | 16 | -0.961 | 0.834 | 0.834 |
| Default Mode | Post | 16 | 16 | -0.722 | 0.765 | 0.823 |
| Dorsal Attention | tFUS | 16 | 16 | 1.036 | 0.155 | 0.198 |
| Dorsal Attention | Post | 16 | 16 | 1.103 | 0.138 | 0.193 |
| Limbic | tFUS | 16 | 16 | 1.345 | 0.093 | 0.163 |
| Limbic | Post | 16 | 16 | 1.445 | 0.078 | 0.163 |
| **Salience** | **tFUS** | **16** | **16** | **2.042** | **0.026** | **0.077** |
| **Salience** | **Post** | **16** | **16** | **1.958** | **0.028** | **0.077** |
| Somatomotor | tFUS | 16 | 16 | 1.413 | 0.085 | 0.163 |
| **Somatomotor** | **Post** | **16** | **16** | **2.328** | **0.014** | **0.066** |

Network (DMN), whereas in *Active*, sgACC exhibited distributed increases to multiple non-DMN networks during and after sonication (Fig. 4). The Cognitive Control network showed the largest Active–Sham differences (ΔFC = +0.19 at tFUS and +0.17 at Post; nominal *p* = 0.0024 and 0.0040, FDR-adjusted *q* = 0.028 for both; Table 3). Positive, albeit smaller, Active–Sham differences were also observed for Visual (+0.042 at tFUS; +0.071 at Post), Somatomotor (+0.096 at tFUS; +0.174 at Post; *q*_FDR_ = 0.066), Salience (+0.131 at tFUS; +0.154 at Post; *q*_FDR_ = 0.077), Dorsal Attention (+0.060 at tFUS; +0.078 at Post), and Limbic (+0.088 at tFUS; +0.093 at Post) networks. In contrast, the DMN showed the opposite polarity (Active–Sham ΔFC = −0.057 at tFUS; −0.039 at Post).

**Fig. 4.**
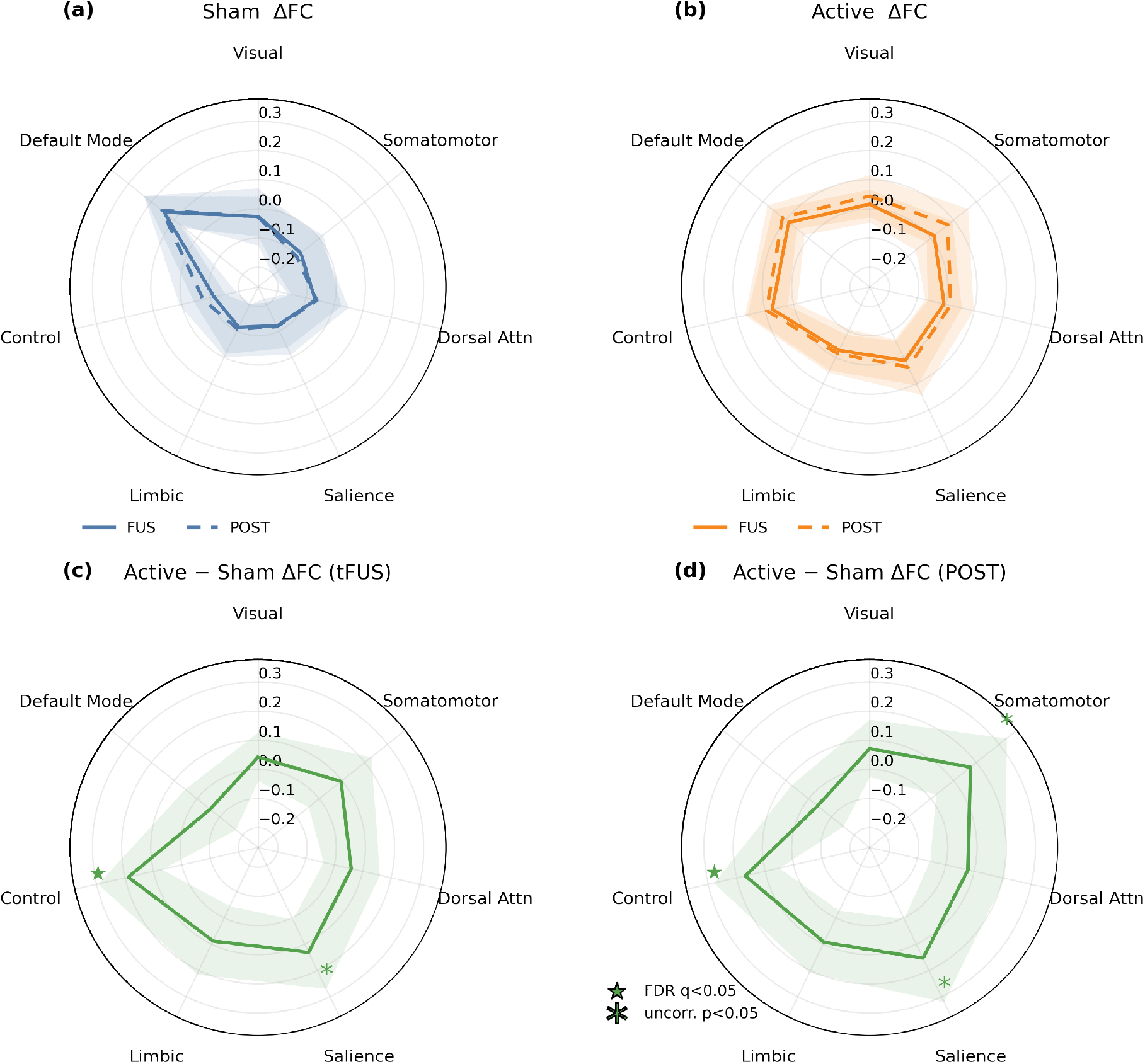
Network specificity of the sgACC connectivity increase under Active tFUS. (a) *Sham sessions:* baseline-referenced changes (ΔFC) in sgACC coupling with the Yeo-7 networks indicate that the sgACC’s connectivity with the Default Mode Network (DMN) increases over the course of the 15 minute recording (`tFUS`: solid; `Post`: dashed). (b) *Active tFUS:* In contrast, ΔFC exhibits a more uniform pattern across the 7 canonical networks, with the largest increase observed at the Cognitive Control network. (c–d) *Active–Sham differences in* Δ*FC* shown at `tFUS`(c) and `Post`(d). Symbols mark directional tests (*H*_1_: ΔFC_active_ *>* ΔFC_sham_) corrected across networks using FDR: filled star ⋆, *q <* 0.05; asterisk *, uncorrected *p <* 0.05. At `tFUS`, the Cognitive Control network’s coupling with the sgACC is significantly positive (FDR *q* = 0.028), with the Salience network showing a positive trend (uncorr. *p* = 0.026, *q* = 0.077). At `Post`, the Cognitive Control network’s coupling remains elevated (*q* = 0.028), while the Salience and Somatomotor networks exhibit trend-level, positive differences (Salience *q* = 0.077; Somatomotor *q* = 0.066). This data is consistent with a redistribution of sgACC coupling from the DMN toward task-positive networks (Control, Salience, Somatomotor) under Active stimulation.

### Examining the influence o f d istance-to-target on FC changes

Due to individual differences in head geometry, the distance from the transducer to the sgACC varied across our cohort (58 *±* 5 mm). The transducer’s fixed beampattern meant that the intensity of stimulation at the target also varied between subjects, and we asked whether this could explain inter-individual variability in observed FC changes. We thus measured the scalp–to–sgACC distance for each subject, and then correlated the result with the FC change using a mixedeffects model (Table 4).

**Table 4.** Moderation of tFUS effects on sgACC connectivity by scalp-to-sgACC distance. Fixed effects included time window (reference = Pre), condition (reference = Sham), z-scored scalp–to–sgACC distance (distance_*z*_; one SD ≈ 5 mm across participants), all two-way interactions (time×condition, time×distance, condition×distance), and the three-way time×condition×distance interaction. A subject-specific random intercept modeled baseline differences. Models were fit by maximum likelihood; Wald *z*-tests (two-sided, *α* = 0.05) are reported for fixed effects. Positive coefficients indicate higher sgACC–whole-brain FC. The Post×Active term corresponds to the difference-in-differences contrast (Active vs. Sham in the Pre→Post change), and the Post×Active×distance_*z*_ term tests whether that effect varies with distance.

| Effect | Estimate | SE | $z$ | $p$ | 95% CI |
| --- | --- | --- | --- | --- | --- |
| Intercept (Sham, Pre) | 0.121 | 0.019 | 6.242 | < 0.001 | [0.083, 0.159] |
| Time: tFUS vs. Pre (Sham) | -0.020 | 0.025 | -0.814 | 0.416 | [-0.068, 0.028] |
| Time: Post vs. Pre (Sham) | -0.011 | 0.025 | -0.451 | 0.652 | [-0.059, 0.037] |
| Condition: Active vs. Sham (at Pre) | -0.061 | 0.025 | -2.485 | 0.013 | [-0.109, -0.013] |
| Time × Condition: (tFUS vs. Pre) × (Active vs. Sham) | 0.060 | 0.035 | 1.731 | 0.083 | [-0.008, 0.128] |
| **Time × Condition: (Post vs. Pre) × (Active vs. Sham)** | **0.079** | **0.035** | **2.261** | **0.024** | **[0.010, 0.147]** |
| Distance ($\text{distance}_z$) | 0.014 | 0.019 | 0.730 | 0.466 | [-0.024, 0.052] |
| Time × Distance: (tFUS vs. Pre) × $\text{distance}_z$ | 0.012 | 0.025 | 0.504 | 0.614 | [-0.036, 0.061] |
| Time × Distance: (post vs. Pre) × $\text{distance}_z$ | 0.016 | 0.025 | 0.639 | 0.523 | [-0.032, 0.064] |
| Condition × Distance: (Active vs. Sham) × $\text{distance}_z$ | -0.012 | 0.025 | -0.501 | 0.617 | [-0.060, 0.036] |
| Time × Condition × Distance: (tFUS vs. Pre) × (Active vs. Sham) × $\text{distance}_z$ | -0.018 | 0.035 | -0.520 | 0.603 | [-0.086, 0.050] |
| Time × Condition × Distance: (Post vs. Pre) × (Active vs. Sham) × $\text{distance}_z$ | -0.030 | 0.035 | -0.872 | 0.383 | [-0.098, 0.038] |
| *Derived contrast (distance-slope difference at Post, Active–Sham):* |  |  |  |  |  |
| $\lambda + \kappa_{\text{post}}$ (per 1 SD) | -0.043 | 0.025 | -1.73 | 0.083 | [-0.092, 0.006] |

*Notes.* Reference levels: time = Pre; condition = Sham. $\text{distance}_z$ is z-scored (distance mean = 58 mm; SD = 5 mm), so coefficients on $\text{distance}_z$ and any interaction thereof are interpreted *per 1 SD of distance*. The key time × condition effect at Post remains significant after including distance (estimate = 0.079, $p = 0.024$ ), indicating robustness of the main stimulation effect to anatomical distance.

In the *Sham* condition, the slope of FC per 1 SD of distance was slightly positive (Pre: +0.014; tFUS: +0.027; Post: +0.030). In the *Active* condition, the slope was near zero to negative (Pre: +0.002; tFUS: −0.004; Post: −0.013). The derived distance-slope difference in the Post time window (Active − Sham) was −0.043 per 1 SD (=−0.074 per 10 mm, SE = 0.025, *z* = 1.73, *p* = 0.083), indicating a *trend* toward attenuation of the FC increase with greater distance-to-target. Importantly, the principal time × condition contrast in the Post time window remained significant after including distance (*β* = +0.079, *p* = 0.024), demonstrating that the main stimulation effect is robust to anatomical distance. Figure 5 depicts the relationship between scalp–to–sgACC distance and change in FC during (a) and after (b) tFUS. Interestingly, application of the Locally Estimated Scatterplot Smoothing (LOESS) technique is suggestive of an inverted U-shaped curve, potentially hinting at the maximization of effects for subjects whose sgACC coincides with the transducer’s focus.

**Fig. 5.**
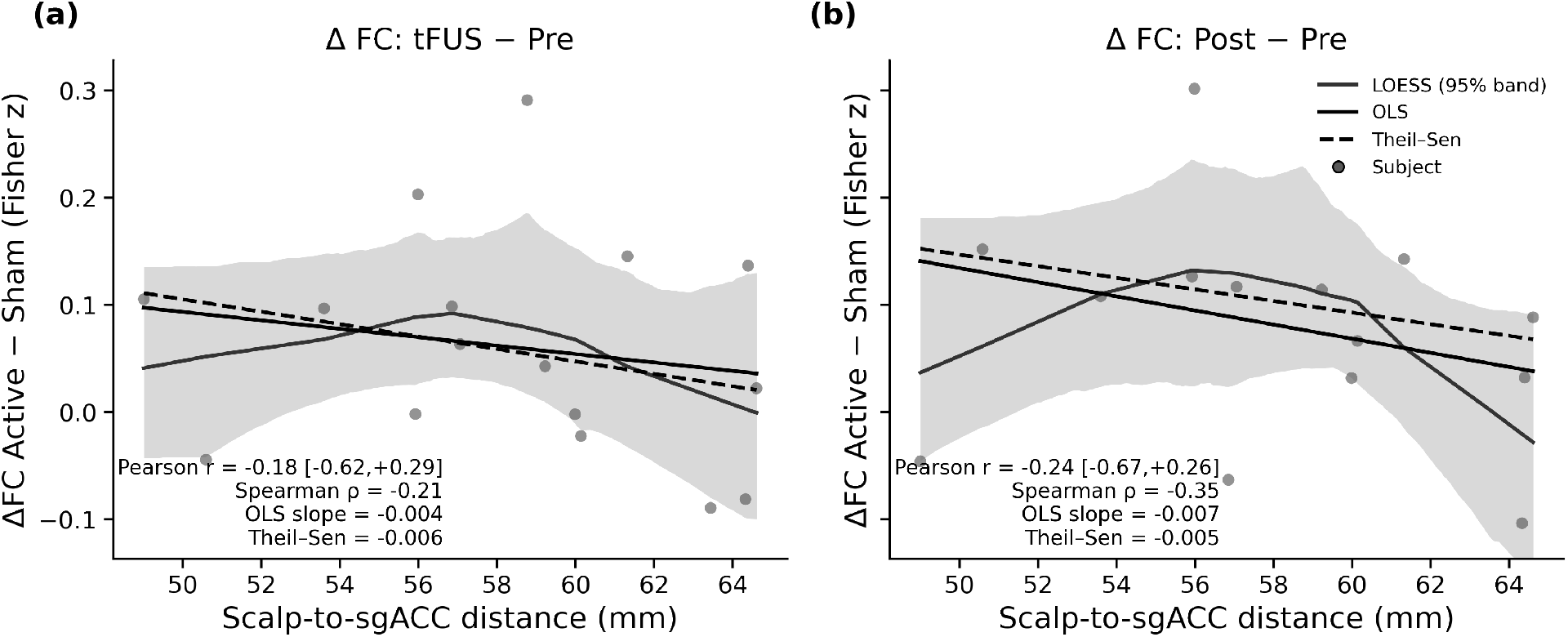
Scalp–to–sgACC distance versus change in connectivity. Panels show subject-level scatter of distance (mm) against ΔFC (Fisher *z* units) for the (a) tFUS and (b) Post-tFUS windows. Each panel overlays a LOESS smoother with a bootstrapped 95% confidence interval (2,000 resamples), a least-squares regression line (solid), and a Theil–Sen robust regression line (dashed). The Post-tFUS panel exhibits a modest inverse trend, visually consistent with a dose-attenuation account; however, the moderation of the Post-tFUS effect by distance falls short of significance in the mixed-effects analysis (*p* = 0.083).

## Discussion

### Summary and interpretation

At a high level, our data show that brief, low-intensity tFUS to the sgACC increases its functional coupling with the rest of the brain in the period *after* sonication, relative to a sham control (e.g., Pre→Post Active–Sham Δ: *β* = 0.079, 95% CI [0.006, 0.151], *p* = 0.033). This effect emerged in a conservative, subject-level mixed-effects model that (i) respects the repeated-measures design, (ii) avoids edge-wise pseudoreplication, and (iii) does not rely on equal baselines, instead estimating a difference-in-differences (DiD) effect between Active and Sham sessions. We did not detect a comparably strong differential effect *during* sonication, suggesting a delayed or accumulating influence of stimulation rather than an acute “online” change.

Analysis of the sgACC’s connectivity with canonical brain networks provides insight into the nature of this accumulating effect. Over the course of the Sham session, sgACC’s connectivity with the DMN increased over time – consistent with the tendency for unconstrained rest to drift toward mind-wandering and internally oriented cognition (48–50). In contrast, Active tFUS did not enhance the sgACC’s coupling with the DMN; instead, its connectivity increases with *non-DMN* systems, most prominently the frontoparietal Cognitive Control network and, to a lesser extent, Salience and Somatomotor networks. The Cognitive Control network is established as a flexible hub supporting goal-directed regulation and cognitive reappraisal (47, 51). The disassociation found here – increasing DMN coupling absent stimulation versus enhanced integration with non-DMN networks during Active sessions —-argues that tFUS is not bluntly amplifying the sonicated region’s activity. Rather, it appears to bias sgACC activity toward circuits implicated in top-down regulation and away from those supporting self-referential processing. Relative to Sham, sgACC-DMN connectivity actually decreased over time, which is interesting in light of the evidence that sgACC-DMN hyperconnectivity tracks rumination and symptom burden in depression (52).

A dynamic systems interpretation of our findings is that tFUS biases ongoing dynamics toward distinct regimes. Restingstate activity naturally explores a landscape in which DMN-dominated states are common; a small, focal perturbation may inject sufficient exogenous drive to nudge the system toward alternative basins of attraction that prioritize control and monitoring. Although we did not measure metabolism, this account harmonizes qualitatively with longstanding observations of sgACC metabolic abnormalities in mood disorders and their modulation with successful intervention (53, 54). If tFUS can transiently reweight sgACC interactions toward control-related circuits, it may counteract the spontaneous drift into DMN-dominant modes that accompany passive rest. This is necessarily a mechanistic hypothesis (i.e., not a conclusion) but it generates concrete predictions for future multimodal work (e.g., Magnetic Resonance Spectroscopy or Positron Emission Tomography alongside tFUS).

A critical aspect of our findings is the enhancement o f FC *following* stimulation. This aligns with prior reports of prolonged tFUS effects in humans and nonhuman primates (12, 55, 56) and is consistent with processes that outlast the sonication epoch. Multiple biophysical pathways have been proposed for ultrasound–brain interactions, including modulation of mechanosensitive ion channels (57), changes to effective membrane mechanics (20), and astrocytic contributions (58). The present BOLD–FC measures cannot adjudicate among these mechanisms. The slow, post-sonication increase observed here suggests a delayed or accumulating influence of stimulation on network organization. We therefore view the persistence of the effect as hypothesis-generating evidence for longer-timescale neuromodulatory or plasticity-consistent processes.

In order to rule out the possibility that baseline connectivity differences between conditions spuriously led to the observed post-sonication connectivity enhancement, we formally analyzed the relationship between the “starting point” of each session and the subsequent change in connectivity. Under a pure regression-to-the-mean account, both conditions should exhibit a similar inverse relationship between baseline FC and the subsequent change. Instead, we observe a robust negative slope in Sham but a mildly *positive* slope in Active, with their difference exhibiting statistical significance (Sham: slope = −0.78, CI [−1.37, −0.18]; Active: slope = +0.21, CI [−0.58, 0.99]). This is inconsistent with a regression-to-the-mean explanation, and indicates that active tFUS alters the baseline–change relationship itself.

Modeling scalp-to-sgACC distance as a continuous moderator yielded a directionally consistent, but trend-level, attenuation of the connectivity enhancement with increasing distance (*p* = 0.083). The local regression analysis hints at an inverted-U shape between distance and effect size (Fig 5). This pattern accords with a dose-attenuation hypothesis, where subjects whose sgACC is closest to the beam focus experience the largest increase in FC. Nevertheless, in our *n* = 16 sample cohort, the post-stimulation effect persists after adjusting for distance, indicating that distance alone does not account for the observed FC changes. Future work incorporating subject-specific acoustic modeling (skull thickness and density), improved targeting uncertainty estimates, and direct intensity proxies may sharpen the sensitivity of distance (or dose) as a moderator.

### Limitations

Our study is not without limitations. Primarily, our modest sample size likely limited our ability to resolve significant effects during sonication, as well as sgACC coupling increases with non-DMN networks other than the Cognitive Control network. In particular, the sgACC-Salience and sgACC-Somatomotor connectivity increase (Table 3) did not survive FDR but showed uncorrected trends (e.g., Salience q=0.08; Somatomotor q=0.07). Secondly, the BOLD signal is challenging to interpret due to the complex biophysics that give rise to its generation (59). In particular, neurovascular coupling is largely agnostic to excitatory versus inhibitory neurotransmission (60), meaning that it is difficult to infer the effect of tFUS on cortical excitability and the overall direction of the change in neural activity. Our employment of FC as the dependent variable – motivated by its well-established association with brain state – further exacerbates this problem, as an increase in connectivity does not imply increased activation of the regions comprising the FC measurement. Alternative forms of MRI such as Magnetic Resonance Spectroscopy (MRS) (12) could permit a clearer understanding of the relative effects of tFUS on excitation and inhibition. Similarly, the utilization of concurrent electrophysiological recordings (7, 61–64) may afford more direct insight into how sonication modulates neural activity. Finally, our study employed a cohort of healthy adult subjects lacking a clinical diagnosis of major depression or other mood disorders associated with impaired sgACC function. As such, it is unclear whether the modulation of sgACC connectivity found here translates to brains whose activity is dysregulated by an affective disorder. Future studies investigating tFUS in clinical populations with aberrant sgACC dynamics are motivated by the present study.

## Conclusion

Brief, low–intensity tFUS to the sgACC produced a sham-controlled, post-sonication increase in whole brain connectivity and a selective redistribution toward control- and salience-related networks, diverging from the DMN drift observed during Sham. These effects were not attributable to baseline differences or simple regression-to-the-mean and showed only trend-level moderation by scalp–target distance, motivating future work that integrates individualized dosing models and multimodal assays to clarify mechanism and translational potential.

## ACKNOWLEDGEMENTS

The authors would like thank Ahmed Duke Shereen at the CUNY Advanced Science Research Center for his help in setting up the combined tFUS-fMRI recordings.

